# Mobile brain imaging in butoh dancers: from rehearsals to public performance

**DOI:** 10.1101/2023.02.26.530087

**Authors:** Constantina Theofanopoulou, Sadye Paez, Derek Huber, Eric Todd, Mauricio A. Ramírez-Moreno, Badie Khaleghian, Alberto Muñoz Sánchez, Leah Barceló, Vangeline Gand, José L. Contreras-Vidal

## Abstract

Dissecting the neurobiology of dance would shed light on a complex, yet ubiquitous, form of human communication. In this experiment, we sought to study, via mobile electroencephalography (EEG), the brain activity of five experienced dancers while dancing butoh, a postmodern dance that originated in Japan. We report the experimental design, methods, and practical execution of a highly interdisciplinary project that required the collaboration of dancers, engineers, neuroscientists, musicians, and multimedia artists, among others. We explain in detail how we technically validated all our EEG procedures (e.g., via impedance value monitoring) and how we minimized potential artifacts in our recordings (e.g., via electrooculography and inertial measurement units). We also describe the engineering details and hardware that enabled us to achieve synchronization between signals recorded in different sampling frequencies, and a signal preprocessing and denoising pipeline that we have used to re-sample our data and remove power line noise. As our experiment culminated in a live performance, where we generated a real-time visualization of the dancers’ interbrain synchrony on a screen via an artistic brain-computer interface, we outline all the methodology (e.g., filtering, time-windows, equation) we used for online bispectrum estimations. We also share all the raw EEG data and codes we used in our recordings. We, lastly, describe how we envision that the data could be used to address several hypotheses, such as that of interbrain synchrony or the motor theory of vocal learning. Being, to our knowledge, the first study to report synchronous and simultaneous recording from five dancers, we expect that our findings will inform future art-science collaborations, as well as dance-movement therapies.

## Introduction

### Background information on the brain architecture of dance

In the past two decades, there has been a mounting interest in identifying the neural underpinnings of artistic expression, and of dance, in particular. The first endeavors towards this direction have focused on studying the brain responses during dance observation, namely while dancers, or non-dancers, perceive videos of dance movements of themselves or others. Brain perception signals have been studied for a variety of types of dance, including but not limited to jazz (1), ballet (2,3), and tango (4), using either electroencephalography (EEG) (1,3,4) or functional Magnetic Resonance Imaging (fMRI) (2). Overall, their findings underscore the power of both techniques to capture signatures subserving differences in an array of dance perception settings (e.g., dance perception by expert dancers vs. nondancers).

Identifying the neural basis of dance performance (i.e., production of dance movements) has been challenging, considering the limitations of neuroimaging techniques that render natural movement in space impractical. Still, researchers have come up with creative ideas to address this question. For instance, Brown *et al*. (5) used an inclined surface in front of the leg room of a Positron Emission Tomography (PET) scanner, where amateur dancers performed small-scale, cyclically repeated leg tango steps while in a supine position. The same group used fMRI to study bimanual partnered movements, with the experimenter sitting next to the lying subject holding hands, and alternating between “leading” and “following” joint movements, similar to those used in tango or salsa (6). In turn, mobile EEG techniques, complemented with motion sensing, have enabled researchers to study brain activity while subjects are dancing freely in the space with the EEG caps on. For example, mobile EEG studies on Laban Movement Analysis (LMA) dancing (7) demonstrated the feasibility of classifying specific movements and LABAN effort qualities from specific EEG signals, and proposed a framework for eliminating motion artifacts from dance analysis. EEG has also been proven effective in picking up not only sex-specific effects during thinking of jazz dancing but also sex-independent effects during physically dancing jazz (8).

It is in this context that we decided to study via mobile EEG the brain activity of five experienced dancers while dancing butoh, recorded simultaneously and synchronously (a process known as hyperscanning), a type of dance and number of dancers that have not been studied thus far, in our knowledge. Moreover, this art-science collaboration allowed us to monitor the creative process through EEG recordings during rehearsals culminating in a theater performance in front of an audience. In this paper, we aim to explain the background, design, neuroengineering methods, and technical validation of our EEG, EOG (electrooculography), and IMU (inertial measurement units) procedures. We report in detail the methodology (e.g., bispectrum estimation, filtering, time-windows, equation) that allowed us to generate a real-time visualization of the dancers’ interbrain synchrony on a screen, while we also propose and demonstrate a signal preprocessing and denoising pipeline that can be used for future offline analyses. Importantly, we share all the raw data and code that resulted from this experiment. Lastly, we discuss how we envision that this interdisciplinary work can inform future art-science collaborations, understanding the brain “in action and context”, and therapeutic practices, using dance as a therapeutic modality for improving wellness and motor deficits.

### Background information on butoh

Butoh is a Japanese avant-garde dance originated by Tatsumi Hijikata and Kazuo Ohno at the height of the counterculture movement in Japan in 1959 (9,10). Although its definition may vary, butoh has been described as a type of dance that allows the exteriorization of bodily reactions, otherwise suppressed in social settings, such as spasms, involuntary jerks, tremor, facial or bodily distortions, falling down, stamping, and rolling on the floor (10). Unlike in other dance styles (e.g., ballet), butoh dancers do not pursue high jumping or fast spinning, they rather focus on their breath and subtle body reactions (10). Butoh has also been seen as a “meditational dance”, due to being a contemplative movement practice that includes deep relaxation and meditative calmness (9). In this, it is similar to other meditative practices, such as Tai Chi or yoga, although, according to Kasai (9), meditation does not picture the essence of butoh as a whole, since the calmness can be interrupted by explosive movements.

These descriptions considered, there are several characteristics that make butoh fall out of the narrow and Western definition of dance (9), while it still taps into common processes employed in a variety of dances. One example is that butoh dancers must be able to execute movements with a modified use of vision; visual stimuli are often shut down, thus the dancer learns to enhance other senses and focus their receptivity to sound stimuli. Another example is that movements in butoh do not tend to be entrained to a periodic metered rhythm, as is the case for most Western dances. Still, the dancers follow sounds and musical cues in the same way dancers in ballet, contemporary dance, tango, or hiphop follow the beat. In the choreography performed for this experiment, half of the soundscape was organized rhythmically, and oftentimes, the dancers were counting, much like in other dance forms. Another section included nature sounds, where the dancers followed sound cues (e.g., pouring rain, bleating of a goat, owl hoot) to inform their movements. There were even parts where the dancers were using the intensity/loudness of the sound as a cue to guide their movements. Thus, while the music and choreography may not be organized in a traditional rhythmic manner, the dancers entrain their movements to other aspects of the sound.

Universal aspects of dance in butoh that are similar to Western styles of dance are, for example, that dancers may move in synchrony and coordinate with other dancers; that they engage in turn-taking to execute movements the one after the other; that they need to synchronize movements between different body parts; that they execute a choreography with self-correction in real-time, while integrating spatial and temporal parameters (i.e., proprioception). When it comes to learning butoh or other choreographies, they all require basic mechanisms of motor, auditory, and sequence learning, as well as short- and long-term memory to learn a sequence of movements in sync with musical or sound cues.

Overall, butoh was considered ideal for this experiment, for reasons that include but are not limited to the following: a) the slow movements that our choreography involved greatly diminished the probability of different motion artifacts; b) the parts of the choreography where the dancers’ eyes are closed allowed us to control for activity in the visual system; c) the motor entrainment to acoustic cues and features, in specific points of the choreography, can be used for analysis around specific acoustic-hallmarks that give greater specificity compared to a continuous periodic rhythm; and d) the choreography was structured so that the movements between the dancers were synchronized only in some parts, leaving space for analysis for both interbrain synchrony during the same and different motor execution, while the acoustic stimuli were controlled for.

## Results

Mobile EEG recordings allow us to study the neural dynamics of dance, individually or in groups, with exquisite temporal resolution (millisecond range) and in ecological settings (e.g., a theater or dance studio) that are not possible to test with other techniques, such as with fMRI, where the subject is constrained to lay down within the confines of the bore of a scanner in a neuroimaging facility. Still, even with mobile EEG recording, there are important technical challenges that we had to overcome, so we present here how we validated technically our EEG, EOG, IMU, and online data synchronization procedures. We also propose a signal preprocessing and denoising pipeline that could be used to analyze our data offline, by showcasing how a fraction of our data looks before and after going through the pipeline. We, lastly, share our raw data and code, both for the live recordings and the live visualization of brain activity on a screen, via an artistic brain-computer interface (BCI) while the dancers were dancing.

### Technical validation of EEG, EOG, and IMU procedures

To record the dancers’ brain activity via EEG, while controlling simultaneously for eye movements via electrooculography (EOG), both at a sampling frequency of 1000□Hz, and head/neck movements via inertial measurement units (IMU), at a sampling rate of 128 Hz, we used two different systems: one 128-channel EEG system split into four 32-channel systems with a Wi-Fi transmitter, and one 32-channel system with Bluetooth communications (see **Methods** for equipment details). We distributed the electrodes following the international 10–20 system (**Fig. 1a**) with slight modifications that allowed for some channels to be used for EOG. Specifically, channels TPO9 and TP10 were removed from the cap and placed on the right and left temples, respectively, to record horizontal eye movement, whereas channels PO9 and PO10 were placed above and below the right eye, respectively, to record vertical eye movements (**Fig. 1a, b**).

**Figure 1:**
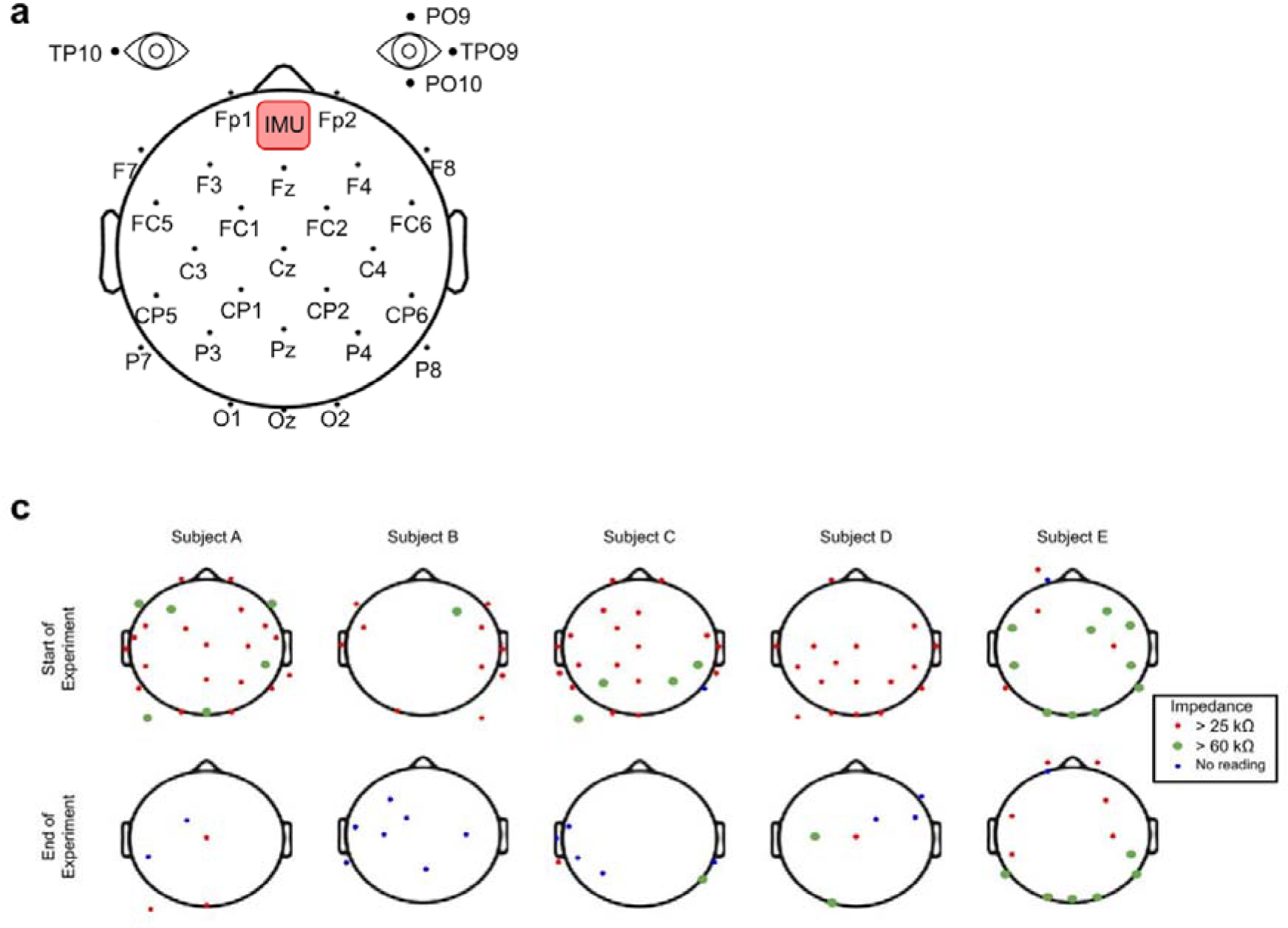
EEG, EOG, IMU locations and impedance values. **a**, EEG-channel montage according to a modified 10-20 system, with ground and reference electrodes placed on the earlobes, channels TPO9 and TP10 on the right and left temples, channels PO9 and PO10 above and below the right eye, and IMU Opal sensors on the forehead. **b**, Close-up image of active EEG electrodes, and location of EOG and IMUs. **c**, Impedance values (kΩ) of the 32-channel EEG for each of five subjects (A-E) at the beginning and the end of the experiment. Impedance values >60 kΩ are considered low quality, and <25 kΩ, of high quality. Subject E was the only participant who used different EEG equipment, with Bluetooth transmission vs. Wi-Fi. [Figure 1b is not shown on BioRxiv due to it including identifiable images.]

The EEG setup was rigorously prepared to minimize potential artifacts that typically contaminate the raw EEG measurements (11,12). A frequent source of noise in the signal comes from electrode artifacts that can occur when there is a disruption in the contact of the electrode with the scalp. In our experiment, for example, this could be driven by the dancer’s sweat while dancing, something further magnified by possible changes in ambient temperature. In the case of the butoh choreography we tested, EMG artifacts would be expected during movements that include head contact with the floor (e.g., while rolling on the floor). All these artifacts would lead to changes in electrical impedance, and, hence, would affect the quality of the recording. Dancing itself might also generate electromyographic (EMG) artifacts originating from the head and neck musculature recruited during head/face and neck movements, which can impoverish the signal, particularly in frequencies above 12-20 Hz, including beta and gamma waves (13). Lastly, although eye movements and eye blinks also typically contaminate raw EEG recordings, there are several methods already published with working protocols to remove ocular artifacts, both offline and online (14), so this artifact did not pose a novel challenge.

To minimize the types of artifacts we mentioned, we followed a set of procedures: a) we measured each participant’s head circumference to allow for selection of an appropriately sized EEG, which would guarantee the right fit of the EEG cap; b) we used a stretchable netting (e.g., medical grade tubular elastic net dressing) to secure the location of all electrodes on the scalp, as well as the electrode cables that otherwise may pull down electrodes during head movements; c) we applied viscous hypoallergenic conductive electrolyte gel between the electrode tips and the scalp to further secure the electrodes in place and reduce electrode impedance; d) we recorded the experiment in a climate-controlled venue; e) we carefully choreographed a piece that was composed to a great extent of slow movements, which helped to minimize motion artifacts; f) we recorded head movements using a distributed system of inertial measurement units (IMUs) to track the dancers’ head motions, so as to later be able to remove motion artifacts from our EEG measurements (**Fig. 1a, b**); g) we designed i) shock absorbing caps for electrode protection during head contact with the floor and ii) travel pillows for neck protection, which we emptied and customized with zippers, to place in the EEG WiFi or Bluetooth transmitters, instead of attaching the transmitters at the back of the head, where the device would be damaged during head contact with the floor (**Fig. 2a, b**); and h) we recorded EOG signals to be able to later remove ocular artifacts and eye movements off-line (**Fig. 1a, b**). All these protection constructs were tested to protect the equipment and were also individually adjusted for the dancers’ comfort. A period of acclimatization was necessary to allow for the dancer to become familiar with the equipment, and its proposed setup, so as to minimize interference with their dance (**Fig. 2a, b**).

**Figure 2:** Customized head and neck protectors used in the experiment. **a**, Shown are scientists carefully placing the electrode caps, processing units, and Wi-Fi transmitters into a neck pillow we customized with zippers (left and right images), as well as three dancers (right image) with custom caps on, which we used as shock absorbing caps to protect the equipment. **b**, Shown are the dancers while dancing in standing (left image) and lying (right image) positions, with their equipment, and head and neck protectors on. **[Figure 2 is not shown on BioRxiv due to them including identifiable images.]**

To assess potential changes or drifts in impedance values, we recorded impedances twice, at the beginning and the end of the EEG recording, using the Brain Vision Recorder software, which allows for visualization of the topographic position of each electrode with a color-coded display of its impedance value (**Fig. 1c**). According to this software, impedance values >60 kΩ are considered low quality, and <25 kΩ, of high quality. To ensure that our data quality would not be meaningfully reduced by high electrode impedance, before starting the experiment, we strived to maintain all impedance values <60□kΩ for all participants. We achieved such values for all Subjects, except Subject E, where there are some >60 kΩ shown (**Fig. 1c**). This correlates with Subject E being the only participant who used different EEG equipment, with Bluetooth transmission vs. Wi-Fi (**Fig. 1c**).

The reliability of such Bluetooth-based g.tech devices is currently a debated topic, with some reporting interpretable signals in the frequency domain and good signal-to-noise ratio (15), while others report poor impedance values, which they interpret might be due to most g.tech devices being equipped with dry electrodes (metal pin electrodes without gel) (16). To address the latter, we used a hybrid gel-based pin electrode for better electrical conduction, which we hypothesized would keep the best impedance throughout the experiment. Nonetheless, our results portray that, despite our efforts, the g.tech device of Subject E still showed poorer impedance values compared to the BrainAmpDC devices. We assume that the inability to improve impedance when the headset was set up influenced the quality of signal and potentially the results, thus we suggest that a BrainAmpDC or similar device would be most recommended for future experiments similar to the one presented here.

### Data synchronization between different EEG measurement modalities and IMU equipment

In our experiment, the use of different measurement modalities and equipment required synchronization between them at a millisecond range to ensure synchronization among the different data streams. To address this issue, we used hardware synchronization via a custom cable for wired transmission of Transistor-Transistor Logic (TTL) signals between devices. In a TTL circuit, transistors act as electronic switches, controlling the flow of current based on the input signals. In detail, as shown in **Fig. 3**, Brain Products’s ActiCAP headsets for Subjects A-D were wired individually to a MOVE EEG transmitter that wirelessly sent their brain activity data to its corresponding receiver via Wi-Fi (Brain Product’s MOVE wireless system). The MOVE receivers sent, via a fiber optic cable, the data to a USB 2 adapter capable of translating the information to a USB cable, which was then readable to the EEG personal computer hosting the Brain Products’s BrainVision Recorder software. The g.tec Nautilus headset of Subject E was also wired similarly to its respective g.tec transmitter but sent data to the g.tec computer via Bluetooth instead. In all Subjects, IMU sensors -attached to their foreheads- sent data to a receiver, with this data being directly sent, via USB cable, to the BrainVision and the g.tec EEG personal computers.

**Figure 3:**
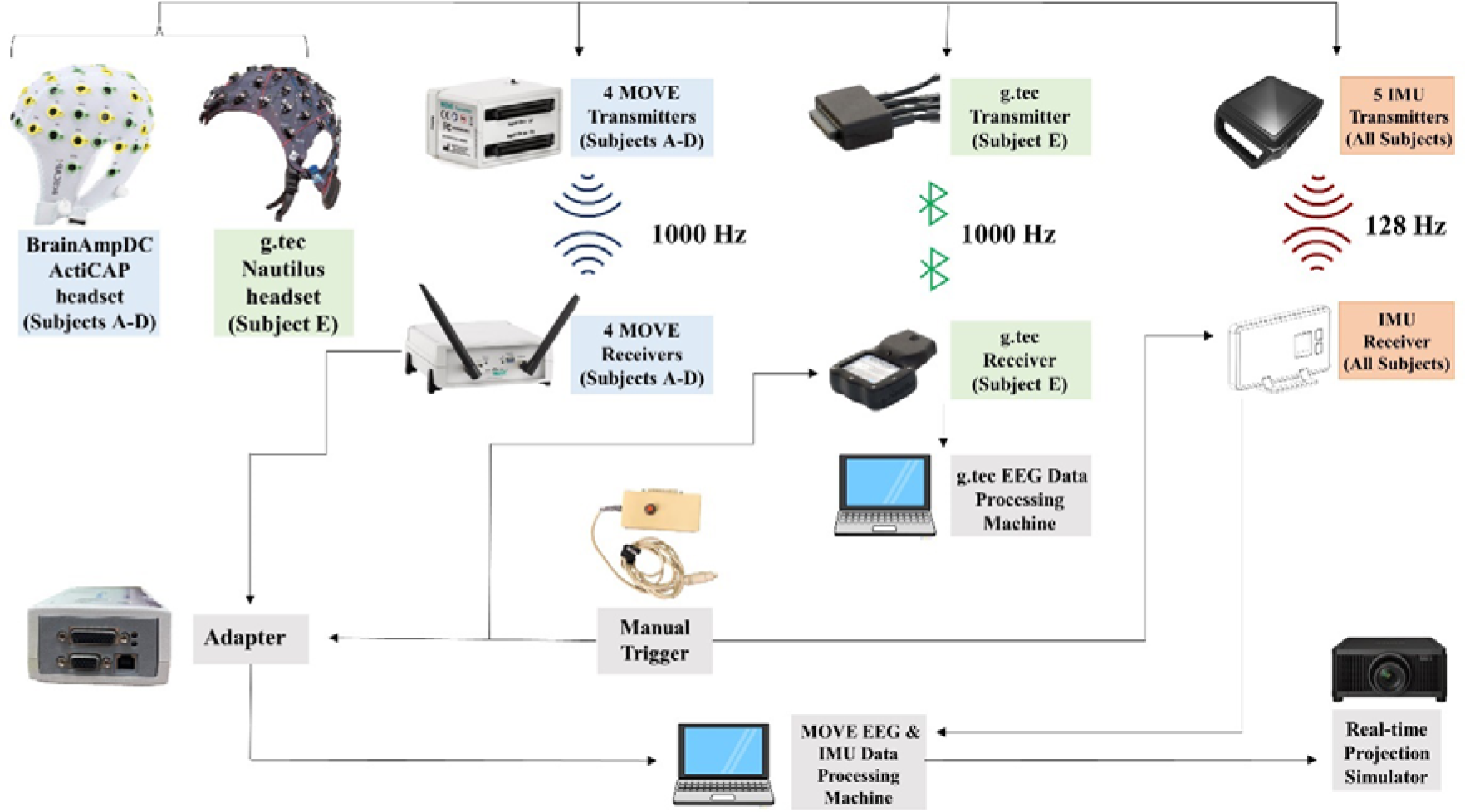
Diagram of the physical configuration of the EEG system in our experimental setup, including hardware elements, connections, and the direction of signal flow. Each box with a name represents an individual device: EEG Wi-Fi-based devices are in blue, EEG Bluetooth-based devices are in green, IMU devices in red, and computer-related devices and other peripherals are in gray. Solid lines connecting boxes represent cable connections between devices. Arrows indicate the direction of signal flow.

Synchronizing the Brain Products and g.tec EEG systems, and IMU sensors required the input of a manual trigger box that had three Transistor-Transistor Logic output connectors (**Fig. 3**): the first one connected to the Access Point antenna to mark IMU data, the second one was connected to the BrainAmpsDC through the USB 2 Adapter to mark WiFi EEG data, and the third one to the Base Station to mark the Bluetooth EEG data. The latter was made possible via a modified dual-pin cable connection. Critical points in the performance and control tasks were marked via the trigger in all three systems at the same point in time. These stored timestamps can be used to align the signals offline for subsequent analysis. **Fig. 4** depicts, as an example, a time-synchronized subset of the recorded raw EOG and EEG signals obtained for 1 second of sitting, speaking, walking, and dancing, with both signals sampled at 1000 Hz. To align signals sampled at different rates, such as those from the IMUs (128 Hz), in the “*Signal preprocessing and denoising pipeline*” section below, we explain how this can be achieved by upsampling the IMU data to match the EEG data by interpolating the timestamps at 1000 Hz (**Fig. 5**).

**Figure 4:**
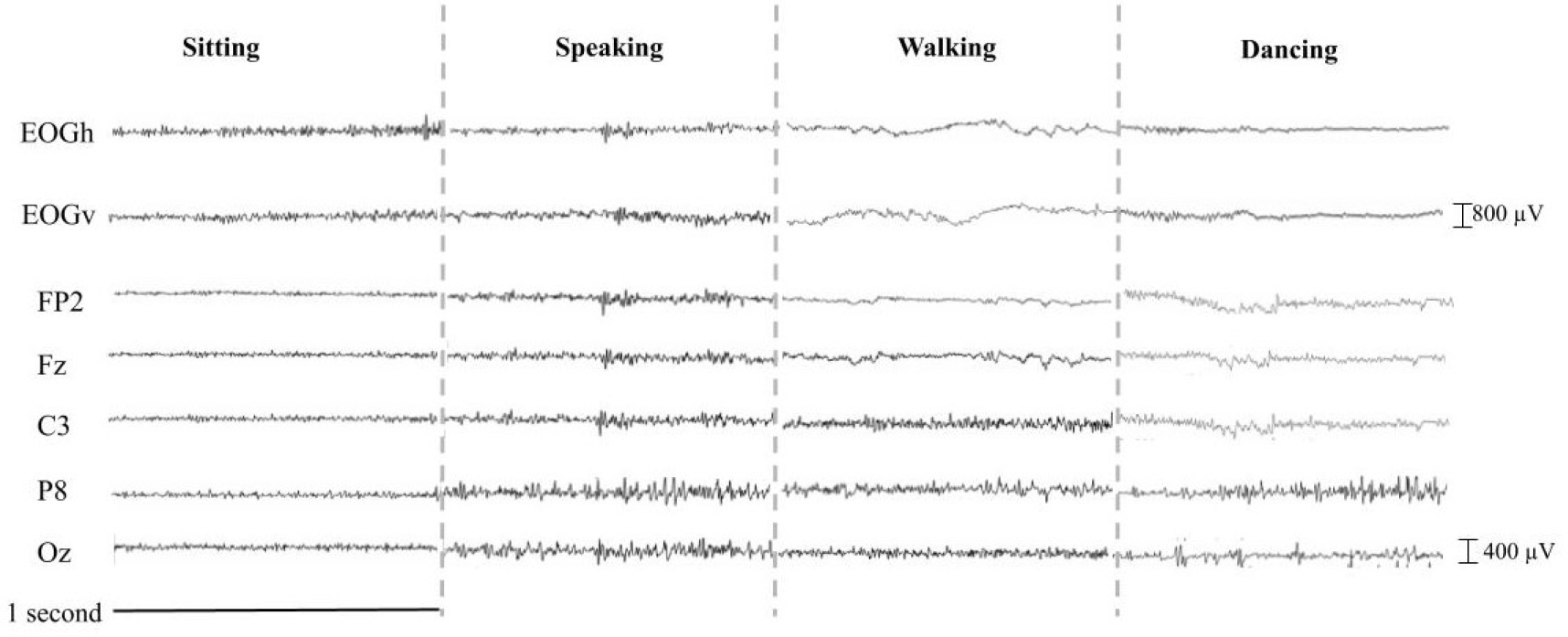
Time-synchronized subset of EOG and EEG during sitting, speaking, walking and dancing conditions. The timeseries EOGh (horizontal) and EOGv (vertical) are computed as bipolar signals for the horizontal and vertical EOG channels, respectively.

**Figure 5:**
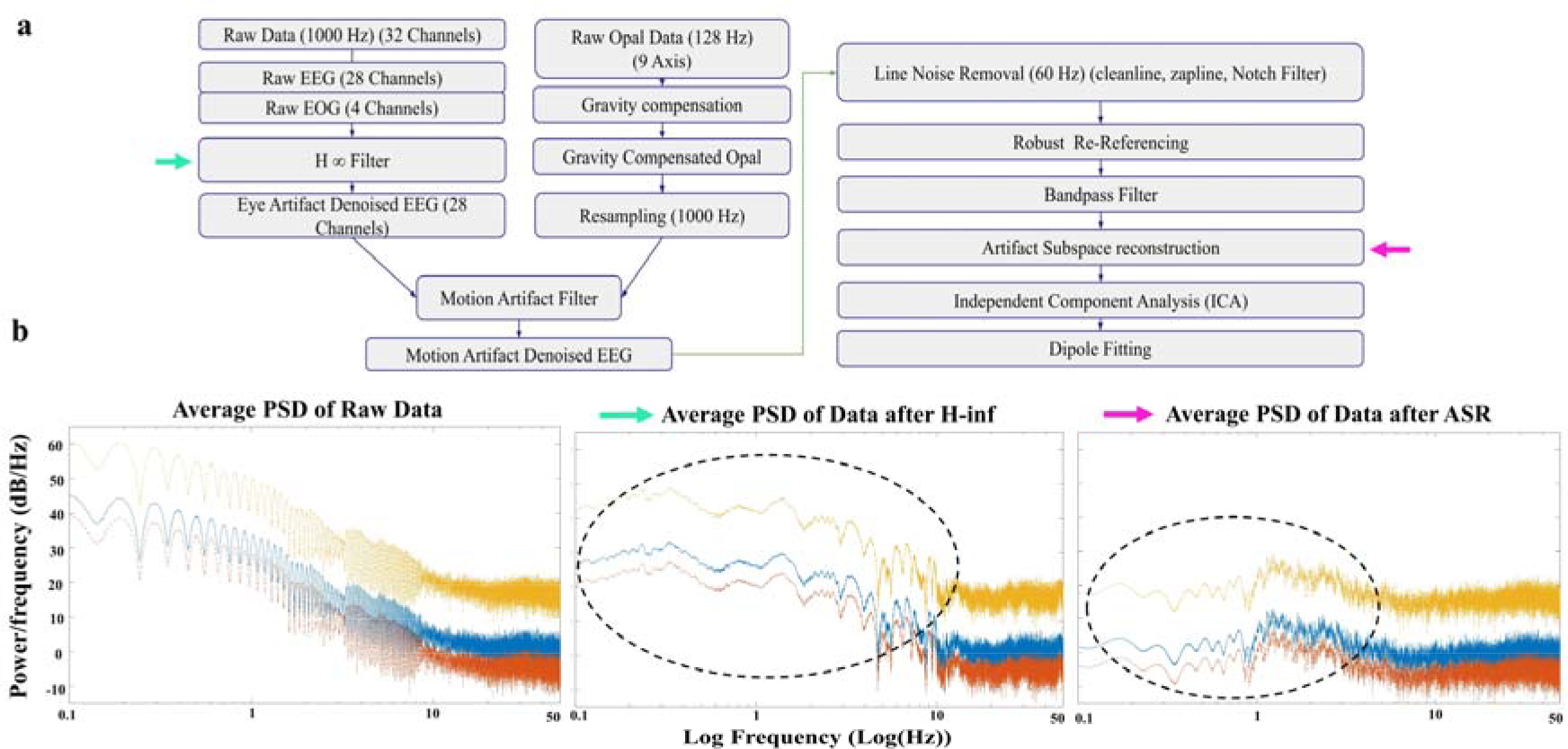
Proposed signal preprocessing and denoising pipeline. **a**, A methodology flowchart showing the basic steps of our proposed pipeline. **b**, Average Power Spectral Density of Subject’s A raw data, data after H-infinity (green arrow) and after the application of ASR (pink arrow) on the interval [0:00 - 20:00]. Dotted circles highlight the main differences between the graphs, e.g., in peak frequencies.

### Signal preprocessing and denoising pipeline

Artifact identification and minimization and noise removal are critical steps in data preprocessing, prior to functional analysis or neural decoding. For subsequent offline analyses using our data, we propose a signal pre-processing pipeline (specific steps shown in **Fig. 5a**; see **Methods** for more details), which ensures the removal of physiological and non-physiological artifacts, including power line noise, as well as achieves a “true” average reference of the signals. Here, we share a demo of this pipeline by using a fraction of our data (**Fig. 5b**). As a first step, since EEG and EOG signals were collected at a sampling rate of 1000 Hz, and the IMU signals at 128 Hz, IMU data was first gravity-compensated and re-sampled to 1000 Hz in order to synchronize it, as well as use it as source noise signal to identify potential motion artifacts in the EEG data in further filtering steps. Following this approach, three-dimensional, gravity-compensated acceleration signals were obtained.

The EEG and EOG raw data were then filtered (H∞ filter (14)) to allow for the removal of eye-related artifacts such as eye blinks, eye motion drifts, and recording biases. These signals were further filtered using an adaptive, non-linear motion artifact removal algorithm in order to preserve the neural content of the EEG signal while increasing the signal-to-noise ratio by removing motion artifacts (17). Motion artifact-denoised EEG data was further cleaned using line noise removal. Following these steps, a robust re-referencing using the PREP pipeline and a band pass filter were applied to further clean the EEG data. These obtained signals were further conditioned using the Artifact Subspace Reconstruction (ASR) algorithm included in EEGLAB. ASR aids in automatically removing transient or large amplitude artifacts that contaminate EEG data. The next step consisted of independent component analysis (ICA), which was utilized to retain brain-related and exclude artifact-related ICs (e.g., residual eye, muscle, electrode popping). In the example comparison we showcase (**Fig. 5b**), we estimated the Average Power Spectral Density of Subject’s A raw data across all 28 EEG channels (1000 Hz) over the first 20 minutes of the choreography with 95% intervals and band-pass filtering between 0.01 Hz and 50Hz, and show how these data compare after the application H-inf and after the application of ASR. Importantly, we highlight the differences in the lower frequencies identified, where after H-inf peaks can be seen at 10 Hz, 25Hz, 30Hz, and near 50Hz), while after ASR, all peaks fall below 20 Hz.

### Real-time interbrain synchronization and EEG-based brain-computer interface visualization

Mobile EEG and brain-computer interface (BCI) techniques allowed us to artistically visualize the interbrain synchrony of the dancers “in action and in context”, during a live dance performance in front of an audience. The artistic design focused on exploring new ways of projecting real-time interactive animated visualizations of EEG data that are both accessible to the lay audience and informative to the audience with a scientific background.

This required that the dancers’ interbrain synchrony be measured in real-time, while dancing butoh, which implies a continuous computation of a synchrony metric across multiple combinations of electrodes of different subjects. The computational load related to this process elevates when the number of electrodes, as well as the number of participants, increase (17). Therefore, to make this online calculation as efficient as possible, we used a Laplacian spatial filter, which filtered the 32 raw channels/Subject into 8 channels (**Methods**). This filter can be thought of as a spatial high-pass filter applied to the data, which attenuates low-spatial-frequency signals that are broadly distributed across the scalp but preserves more localized higher-spatial-frequency signals. As shown in other studies, higher bispectrum magnitudes at given pairs of frequencies reflect non-random interactions, phase coupling (18), and non-linear multi-frequency interactions (19), which have been observed as traces of interbrain synchrony, for example, during teamwork interactions (20). Using these filtered 8 channels/Subject and calculating the bispectrum between dyads across the gamma frequency band (between 30 Hz-50 Hz) yielded blocks of 8 by 8 matrixes which were used for the BCI visualization.

Our decision to use these filtering parameters, and not use further signal denoising, are justified by our aim to run a bispectrum analysis in real-time, where we could not afford more time and space-consuming methods. For example, in offline analyses, more complex and computationally slower filters can be used to remove unwanted artifacts from different sources, such as robust referencing, Artifact Subspace Reconstruction (ASR), and Independent Component Analysis (ICA) before analyzing the data. Additionally, in offline analyses, component space can be used, as opposed to channel (sensor) space, which allows for more accurate localization of the brain areas activated. Lastly, offline analyses give more flexibility in the time-windows and overlaps that can be used to estimate synchrony, such as 2-second windows with 50% overlap or 4 second-windows with 75% overlap, while real-time bispectrum can be estimated using a single time-window and overlap (in this case, 4 second-windows with 75% overlap).

The BCI visualization of the resulting data was inspired by the structure of the music composed for the purpose of the study, the choreography itself, and the concept of interbrain synchrony. Since butoh is often made up of slow movements, and in contrast, the music for the project is active and repetitive, the visualization aspired to find a middle ground between the physical and sonic rhythms. Overall, the visualization had no identifiable regular pulse but flowed freely in terms of texture, movement, color, and spatialization. It was structured in three sections: a) abstract scenic monochromatic images, b) five 3D brains and brain connections portraying interbrain synchrony, and c) five abstract circles producing a colorful texture (**Fig. 6**).

**Figure 6:**
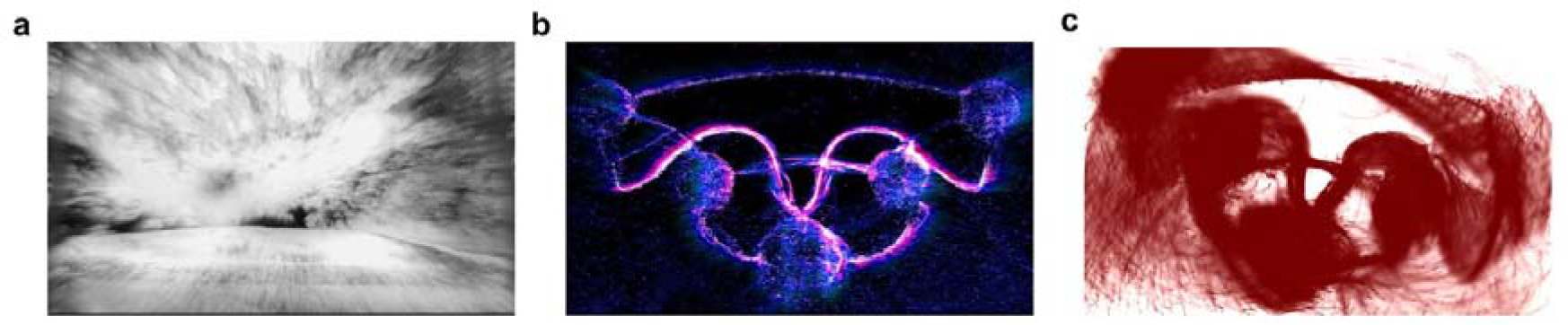
Real-time interbrain synchrony visualization via brain-computer interface. **a**, Abstract scenic monochromatic images, with additive noise, whose amplitude was the result of the leading dancer’s normalized raw EEG data. **b**, Five 3D brains and their connections reflecting real-time brain synchrony. **c**, Five abstract circles showing the interbrain synchrony between the accompanying dancers and the dance leader. In all cases, top images show computer examples from the type of visualization described, and bottom images show real instances of how these visualizations unfolded during a live performance. [Bottom images are not shown on BioRxiv due to them being identifiable images.]

In detail, at the beginning of the music and for the first 20 minutes, the visualization was meant to depict the prevailing sounds of nature with abstract scenic monochromatic images with additive textural visual noise. This noise’s amplitude was the result of mapping normalized raw EEG data of the dance leader (**Fig. 6a**). The second visualization of the five 3D brains and their connections reflected the positions of each dancer on stage to help the audience associate the brain visuals with each dancer. The brain synchrony value was mapped in real-time with the particle flow level between brains, forming a line between them, so that the higher the value of synchrony, the higher the opacity and thickness of the line (**Fig. 6b**). The third visualization of five abstract circles showed the interbrain synchrony between the accompanying dancers and the dance leader. Their interbrain synchrony was mapped to the position of the four circles relative to the central circle, so that the higher the value of interbrain synchrony, the more concentric the circles appeared (**Fig. 6c**). We believe that this visualization was crucial in communicating the essence of this collaboration, right at the intersection of art and science.

## Discussion

In this study, we reported in detail the experimental design, methods, technical validation, and practical execution of a highly interdisciplinary project, which is meant to inform future projects. We specifically listed the best practices and steps that we took to achieve high-quality impedance values and shared our observations and suggestions on the devices that performed the best. We also shared our practices in recording different types of artifacts (via EOG and IMU) to maximize the signal-to-noise ratio. Additionally, we explained how we synchronized different EEG devices, via TTL triggers, and how we ran interbrain synchrony analysis in real time that we then used for the BCI visualization. For future reference, we put forward a signal preprocessing and denoising pipeline that scientists can apply using the raw EEG data and code we shared here. We believe that this data can be used to address different hypotheses. In what follows, we explain several of these hypotheses that we envision.

### Hypotheses to test

#### a) Interbrain synchrony

Dance has been posited to have evolved as a form of interpersonal coordination and social communication, which is based on both imitation (matching of movement) and synchrony (matching of time) skills (21). Different kinds of dances rely on different aspects of interpersonal coordination, including touch, eye gaze, sensory-motor interactions, facial expressions, or even synchronization with other physiological parameters, such as breathing, heartbeat, and sympathetic tone (22). Thus, EEG recording from different dancers, dancing the same choreography simultaneously, is expected to unravel interbrain neural synchrony in the dance aspects that require interpersonal coordination.

Previous experiments (23) where we examined EEG signals of two dancers while dancing a ballet duet showed high interbrain synchrony in the gamma band of visual brain regions (Broadman area 18) of the dancers. Interestingly, the leading dancer also showed interbrain synchrony between her visual (BA18) cortex and her partner’s cognitive (BA31) and premotor/supplementary motor areas (BA6) of the brain. The butoh choreography that we investigated was focused on auditory cues from the music, meaning that the dancers were proceeding from one move to the next in response to specific auditory cues, such as a sound that resembles an owl hooting announcing a specific move. Throughout most of the choreography, the dancers danced with their eyes closed or half-closed with a soft focus (24), something that gives us the unique opportunity to study interbrain synchrony in a type of dance where vision is not expected to be the basis of coordination, reported as the most common form of interbrain synchrony (25). Lastly, since we recorded both the rehearsals and the final performance, it will be possible for us to assay changes in interbrain synchrony as a function of learning, practice, or the scenic context (e.g., presence or absence of an audience).

For offline analyses of interbrain synchrony, we recommended specific signal preprocessing and denoising pipelines, as well as filtering practices, such as robust referencing, Artifact Subspace Reconstruction (ASR), and Independent Component Analysis (ICA). We also suggest that component space, instead of channel space, be used for bispectrum estimations, which allows us to ask questions by zooming in on specific Brodmann areas, as we have previously shown (23). Another interesting practice would be to use differing time windows and overlaps to yield more fluid changes in interbrain synchrony, such as 2-second windows with 50% overlap and 4-second windows with 75% overlap.

#### b) Motor hypothesis of vocal learning

There is a hypothesis (26) that links the evolution of the neural circuit that is responsible for rhythmic body muscle movement (e.g., head, arm, and leg muscles) to the evolution of the neural circuit that is responsible for the movement of the muscles of the vocal organ during vocal communication (e.g., laryngeal muscles in humans). This hypothesis is built on findings (26) showing that in vocal learning birds, all their cerebral nuclei that are devoted to song learning are adjacent to discrete brain areas active during limb and body movements. In other words, the hypothesis states that our ability to move in time with an auditory beat (or, dance, in humans) originated from the neural circuitry for complex vocal learning (or, speech learning, in humans).

This prediction became even more pertinent after the finding that only species that communicate with complex vocalizations (i.e., humans and parrots) are able to dance (i.e., to entrain their body movements to a beat) (27), pointing to a common neural substrate in both abilities. In humans, this hypothesis has not been tested, meaning that no one has compared in the same subjects the neural pathways underlying speech (i.e., laryngeal movements) and dance movements (e.g., rhythmic arm movements), although a cross-studies’ comparison points to an overlap of several of the regions controlling body movements in the primary motor cortex with the regions that control laryngeal movements in the primary motor cortex (28–30). Further, dance has been found to increase network connectivity between the basal ganglia and premotor cortices (31), with both regions being co-activated during speech (32,33).

To address this question, in our control tasks, we had our dancers produce speech and speech-like vocalizations (e.g., Jabberwocky words), as well as other non-speech vocalizations (e.g., sneeze, laughter, yawn), with the aim to compare their EEG patterns during laryngeal movements vs. movements of other body parts. We suggest that for the comparison of these two types of movements, both short (e.g., 1 second) and long (e.g., 4 seconds) time windows would serve different purposes, as long time windows would capture all relevant activity, but might include more artifacts, compared to shorter windows. We also suggest that the best dancing movements for this comparison would be movements that do not require a lot of physical strength, as heavy breathing activates the laryngeal muscles even further. One way to control for the laryngeal movements of breathing would be to subtract the brain activity during resting state, when the subjects were seated and breathing, from both the speaking and dancing activities.

#### c) Butoh vs. other meditative practices

As aforementioned, butoh embraces in its practice contemplation and meditation (9), so it may tap into similar processes as those involved in, for example, seated meditation, such as attention mechanisms that guide concentration. There are also embodied meditative practices, such as yoga and tai-chi, with which butoh might be similar, in that all they include movement in different degrees while maintaining focused attention, and some of them include auditory attention to external stimuli. As such, our butoh EEG recordings offer a comparandum for other meditative practices, where there are already published data on their associated brain activity.

For example, Banquet (34) used spectral analysis of the EEG during transcendental meditation, a method described as a mental repetition of a special sound or mantra, to show in the early 1970s that meditative states can be distinguished from other states of consciousness based on sequential changes in the alpha, theta, and beta waves in relation to their topographical alterations across the scalp. More recently, EEG recording during meditation in Buddhist practitioners revealed self-induced and sustained high-amplitude gamma-band oscillations (35). In a different study, meditation training gave rise to increased theta activity in the frontal midline electrodes, which was sustained even during the resting state following meditation training (36). Xue *et al*. (37) in a similar experiment on short-term meditation training found increased theta (and some alpha) activity in the anterior cingulate cortex and adjacent prefrontal cortex, which correlated with improved performance on tasks of attention, working memory, creativity, and problem solving.

All these studies provide a fertile ground for comparison with the hypothesized meditative aspects of butoh. To make the comparison between butoh and meditation readily possible in our experiment, we included in our control tasks a seated meditation task, which will be compared with the time frames of the butoh choreography that employ contemplative practices. In this way, we will be able to address the question of whether what is happening in the brain during seated meditation bears any resemblance to butoh in the same subjects. Whether the signatures are the same or different, given the evidence showing that meditation-like practices are beneficial in several aspects of human health (38), we hope that our experiment will shed light on the patterns of brain activity underlying these practices.

#### d) Butoh in pregnancy

Throughout the experiment, the pregnancy of one of the butoh dancers gave us the opportunity to study brain activity during butoh dancing in a pregnant woman. To our knowledge, this is the first time to run mobile EEG with a pregnant woman dancing butoh, or dancing, in general. Concerning the safety of dancing during pregnancy, there is published evidence that dancing can actually be beneficial in pregnancy, as long as it does not include lifting other dancers, or high-impact activities such as jumping and back flips (39,40). Considering that, in butoh, such movements are rare to begin with, we deem that this coincidence might uncover butoh’s untapped role as a beneficial practice in pregnancy. For the purpose of the experiment, the choreography was specifically altered to suit the movement abilities and safety of the pregnant woman, and the performance was expressly allowed by a doctor.

Regarding the use of EEG during pregnancy, there are already published reports on the safety of this technology in pregnancy (41). Interestingly, Plamberger *et al*. (41) used a visuospatial attention task, where an auditory cue directed the attention of pregnant and non-pregnant participants either to the left or to the right visual hemifield, where, following a variable time interval, they had to discriminate between a “p” or “q” sound on the cued hemifield. Both non-pregnant and pregnant women showed a decrease in the alpha amplitude in the fronto-parietal network, which correlated positively with accurate discrimination, with no significant differences in the cases of pregnancy vs. non-pregnancy. Since our butoh choreography is based on correctly perceiving auditory cues in the music, it is tempting to hypothesize that an alpha band desynchronization, leading to the expected cue and right after the cue is perceived, could underlie accurate choreography performance, and to study whether there are any differences between the non-pregnant dancers vs. the pregnant dancer.

### Live test for interdisciplinarity

As a collaboration studying butoh in the brain, both the art -butoh- and the science -EEG recording-were equally important towards success (42,43). This live test for interdisciplinarity allowed exploration into understanding the unique opportunities and challenges for such a collaboration, including needs for dancers as athletes and subjects, technical requirements for protecting equipment without inhibiting movement, and for synchronizing brain waves across all five dancers, implications for providing education and working with students, and determination of visual projection based on BCI.

Because this study relied on hyperscanning, the condition of sound was a critical component, not only for live visualization via BCI but also because auditory cues were often the only signals upon which to coordinate motor movements. Thus, any discrepancies or failures in sound quality would greatly impede the dancers’ performance, and potentially relatedly, EEG recordings based on the ability to enter into anticipated parts of the choreography and synchronize with each other. As a result, this would percolate down to the real-time interbrain synchrony calculated and to the BCI-visual projections.

This collaboration further highlighted that bringing various fields together requires clear communication to understand the various needs and expectations of each discipline (44,45). For example, for dancers, who are highly skilled athletes (46), a controlled environment, considering aspects of stage size, noise level, temperature, and other factors that may affect the dancers’ performance, must be considered to minimize stress and maximize their ability to perform. For scientists, it is also critical to factor in human fatigue in EEG recordings; simply recording data, if the question at hand is as specific as that we are asking about butoh dancing, will not suffice, and errors in sound production, unnecessary delays lengthening the time of preparation for study, and any other factors that may impede the dancers’ ability to perform butoh may lead to poorer data outcomes.

Given that the dancers are the subjects of interest, this collaboration also showed that creative solutions may lie in another’s lived experience. One example was that the solution for how to best don the EEG caps, which are sensitive both for capturing brain waves and as a piece of equipment, was found by one of the dancers. Another example comes from a dancer who reported that the control tasks are better to be recorded before any butoh dancing takes place, since the butoh (meditative-like) state may linger after the performance and confound results from control conditions. These situations are great examples of the benefits of this multidisciplinary collaboration, in which the perspectives from experts in different fields came into play and merged into a highly unique project.

Ultimately, seemingly competing interests for data needs must all be considered as all interested parties have their own reporting requirements for meeting funder expectations and for future funding: for dancers, a high-quality video; for scientists, robust data collection; for students, time and attention for hands-on learning. The reader is referred to a recently edited book on *Mobile Brain–Body Imaging and the Neuroscience of Art, Innovation and Creativity* (47) that addresses the challenges and transdisciplinary opportunities for transformational and innovative research and performance at the nexus of art and science enabled by emergent technologies.

### Dance -and butoh-as movement therapy

Dance-movement therapy is the use of creative movement (48) as a healing tool rooted in the inseparable connection between the body and the mind, with concepts of embodiment and attunement affecting human behavior –psychologically, physically, and socially (49). The therapeutic effect of dance is reported as a modality for health and wellness across the lifespan, including for motor development in children in general (50), with Down syndrome (51), cerebral palsy (52), and developmental cerebellar anomalies (53), through the elderly for successful aging, for markers including fitness, functional balance, mobility control (51–53), and cognition (54). Overall, dance-movement therapy and dance leads to psychological health outcomes including decreasing depression and anxiety, increasing quality of life, and expanding interpersonal and cognitive skills (55).

Among patients with Parkinson’s Disease, music, and dance proved to be simple, non-invasive treatment options that promote balance, gait, and cognition (56–59), decrease psychological symptoms, and improve quality of life (60,61). For other conditions, such as schizophrenia and psychotic disorders, many studies tend to have small samples, no randomization, and no adequate control (62). Yet, there is some support that body-centered interventions do alleviate stress, depression, and anxiety as well as facilitate pain reduction in physical and psychological pathologies via a bidirectional pathway between the brain and body (63). As one creative therapy, dance has been shown to be effective for severe mental illnesses such as trauma-related disorders, major depression, and bipolar disorder (64,65).

Because butoh dance is a psychosomatic exploration method (9), this study holds implications for further understanding the healing effects of dance, particularly how dance-movement can be prescribed as a form of therapy by selecting the emotional or physical level of involvement, or dose response, based on the patient’s condition. For example, there may be a relationship between some mental aspects of schizophrenia and butoh performance in terms of the state of consciousness and body-mind vulnerability (66).

## Conclusion

The art-science collaboration that we reported here was a unique, complex, multidisciplinary experiment that required the coordination, managing, and execution of several dancers, engineers, neuroscientists, musicians, multimedia artists, logistic personnel, facilities’ management crew, and students. In addition, securing funding for the travel expenses and artists’ fees was critical to the success of the project. Last but not least, trust and respect for each other were essential to conduct the project in an accelerated timeline. The resulting data, best practices, approach, code, and audiovisuals present a unique opportunity for the scientific and artistic communities to harness the data, knowledge, and lessons learned from this project, to answer novel questions, deploy new algorithms or computational methods, and create new art-science works.

## Methods

### Participants

Five healthy female adults with no history of neurological disorder, or movement difficulties participated in this study. The experimental protocol and informed consent (reviewed and signed by each participant) were approved by the Institutional Review Board (IRB) at the University of Houston. All experiments were performed in accordance with the 45 Code of Federal Regulations (CFR) part 46 (“The Common Rule”), specifically addressing the protection of human study subjects as promulgated by the U.S. Department of Health and Human Services (DHHS).

### Instrumentation & Data Collection

High-density active electrode scalp EEG and electrooculography (EOG) recordings were obtained simultaneously for the five dancers at a sampling frequency of 1000□Hz. For EEG recording, we used two different systems: one 128-channel EEG system (BrainAmpDC with Acticap active electrodes; Brain Products GmbH, Munich, Germany) split into four 32-channel systems with a Wi-Fi transmitter, and one 32-channel system (Nautilus, Gtec medical engineering GmbH, Austria) with Bluetooth communications. Channels TPO9, PO9, PO10, and TP10 were removed from the cap and used for electrooculography (EOG) to capture blinks and eye movements. Channels TPO9 and TP10 were placed on the right and left temples, respectively to record horizontal eye movements, whereas channels PO9 and PO10 were placed above and below the right eye, respectively, to record vertical eye movement. The remaining 28 channels were arranged according to the 10-20 system (Fp1, Fp2, F7, F3, F4, F8, FC5, FC1, FC2, FC6, C3, Cz, C4, CP5, CP1, CP2, CP6, P7, P3, Pz, P4, P8, PO9, O1, Oz, O2, PO10), with ground and reference electrodes placed on the earlobes.

In preparation for the experiment, the dancers were asked to refrain from using products in their hair that may increase the impedance at the scalp/electrode interface (e.g., conditioner, hair gel, etc.). Before donning the cap, the skin on the face around the eyes, the temples, and the earlobes were gently cleaned with alcohol wipes to remove any dirt and skin oils. The cap was aligned on the head such that the FP1 and FP2 were 10% of the distance from the nasion to the union along the midsaggital plane, and electrode Cz was at the vertex of the head. Finally, a light surgical mesh was placed over the electrodes to fix their location on the scalp and to mitigate the potential influence of motion artifacts during the control movement and dancing tasks. We used the Brain Products MOVE system to wirelessly transmit the EEG data to the recording computer. We connected the cap electrodes to a wireless transmitter, which we placed inside a customized neck travel pillow, which we emptied. The WiFi receptor was connected to the BrainVision Recorder computer for storage and processing of the EEG signals.

Head motion signals were recorded using a distributed system of inertial measurement units (IMU; Opals, APDM Wearable Technologies Inc, Portland, OR) to track the dancers’ head motions. The IMU Opal sensors were placed on the forehead of each dancer, acquiring data at a sampling rate of 128 Hz. The data was stored on an onboard micro SD card and simultaneously streamed to a PC for visualization and data monitoring using the Biometrics Analysis Software (Biometrics Ltd, Newport, UK). The recording sites were cleaned with an alcohol solution and allowed to dry. The sensors were then fixed to the skin using a double-sided adhesive tape (designed specifically for Opal sensors, so as not to obstruct electrodes). The signals were calibrated and checked for quality while the subjects stood in a neutral posture with their hands by their side, or while moving their head/neck.

### Experimental Protocol

The experiments were conducted over a series of 4 days, with each day differing in the data collected. The first recording (Day 1: 12/9/22) consisted of a rehearsal only with subject C, where we were able to test the fit of the equipment and several of the technical procedures we followed. On days 2 (2/6/23) and 3 (2/7/23), we recorded a series of control tasks with all subjects (A, B, C, D, and E). Days 4 (2/8/23) and 5 (2/9/23) involved calibration, rehearsal and control tasks with all subjects. Day 5 additionally included the recording of control tasks, calibration, and final performance in front of an audience. All the experiments were conducted at the University of Houston Student Center South Theatre.

Before starting the control tasks, 1 minute of EEG and EOG were recorded to establish a baseline period of brain and muscle activity. Afterwards, the control tasks included a resting state at both the beginning and the end (1 min); walking (2 min); vocalizations (10 minutes) that included a) reading a list of words (“snake, wind, mist, hitch, brief, vent, throat, click, jeeps, mouse”), b) reading a list of nonwords (“brant, pipso, brab, blave, filt, golk, raint, tane, praine, shaty”), c) reading a list of “Jabberwocky” words/sentences (“The blay florped the plenty mogg”; “The Gou twuped the vag all lus rall”; “The heafest dropding deak is rhaph phemes away”), d) producing sentences (“My name is Vangeline. I live in New York. Butoh is great, isn’t it?”), and e) producing volitionally vocalizations that are typically produced spontaneously (laughter, sneezing, and yawning); seated meditation (5 minutes); butoh movements done mechanically, without “dancing” them, without being into the butoh mood, and with no music (7 min); and simple movements, such as raising right/left arms and legs, doing cyclical wrist/ankle movements, tongue protrusions, lip movements (e.g. lip rounding), nostril movements, finger/toe movements, and opening and closing the jaw (completed in sequences of 10 movements for each task; 10 minutes).

The 60-minute butoh choreography the dancers performed was choreographed by Vangeline Gand under a series of specified parameters. These parameters included variations in speed (e.g., no movement, motor preparation, very slow movements not detectable to the eye, slow movements detectable to the eye, pedestrian/walking speed), type of muscle contraction, butoh technique used, choreography type, imagery, gravity (e.g., working against gravity involved muscle tension, while the opposite, muscle relaxation), and emotion.

### Interbrain synchrony

The quantitative measurement of interbrain synchrony that was used for the BCI visualization was achieved by calculating the bispectrum between dancer-dyads of EEG data obtained while performing butoh. Specifically, bispectrum combinations were generated (1 Hz) between the following dyads: subjects C-A, C-B, and C-D. Data from Subject E were excluded because of the possible differences in quality due to using a g.tec device.

Bispectrum was estimated across the EEG recordings using 4-second windows with 75% (one-second) overlap. The bispectrum at each time window was estimated using **Equation 1**:

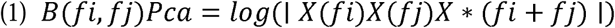

where subscripts Pca are for the data used from Participants “C” and “A”, in this example, and fi the frequency vector for the signal of Pc, and fj the frequency vector for the signal of Pa. X(fi) and X(fj) represent the Fourier transform of window l at frequencies fi and fj respectively. The term X∗(fi+fj) represents the complex conjugate of the Fourier transform of the sum of both frequencies fi and fj (67). The quantity B for bispectrum is obtained by taking the logarithm of the absolute value of the product of the Fourier transforms and their complex conjugates at frequencies fi, fj and fi+fj. Using this method, bispectrum was estimated for all fi = fj, in 50 frequency bins between 1-50 Hz.

The 32 raw channels for each of the 4 subjects (A, B, C, and D) were filtered into 8 channels using a Laplacian spatial filter. Then bispectrum between Subject C and each of the other Subjects for gamma frequency was calculated using the formula (Equation 1), yielding an 8 by 8 matrix comparing each of the 8 channels between two subjects for the three dyads/combinations we tested (subjects C-A, C-B, and C-D).

### Artistic brain-computer interface

The visualization was designed by TouchDesigner (TD, Derivative, Toronto, CA), a visual programming environment aimed at real-time 3D rendering, combined with high-resolution real-time compositing (https://derivative.ca/). MaxMsp, a visual programming language for music and multimedia developed and maintained by San Francisco-based software company Cycling ‘74 was used in this project for data filterings and mathematical operations such as normalizing, scaling, averaging, and calculating min. and max. of input data. Additionally, MaxMsp was used for optimizing the computation and construction of the user interface for the change of the visual scenes.

To establish a mechanism to translate the EEG data to artistic visualization (BCI), the science team and multimedia artist established a communication system between MATLAB (The Mathworks Inc., Natick, MA) and TD. Two data packets were transferred from MATLAB to TD using the networking protocol TCP/IP (https://docs.derivative.ca/index.php?title=TCP/IP_DAT) via a direct Ethernet connection. The first packet of data is the raw EEG data of the dancers with a frequency of 100 Hz. The second packet is the interbrain synchrony with a frequency of 1 Hz. The received data transfers to MaxMsp via Open Sound Control (OSC), a protocol for network communication among computers, sound synthesizers, and other multimedia devices (https://www.cnmat.berkeley.edu/opensoundcontrol). MaxMsp filters/calculates the data and sends them back to TD via OSC.

### Signal preprocessing

EEG signals were pre-processed utilizing MATLAB R2023a (MathWorks, MA), and functions from the open-access toolbox EEGLAB. The EEG and EOG raw data were filtered with the Adaptive Noise Canceling (ANC) H∞ filter. For this step the parameters γ = 1.15, q = 1e-10, and Po = 0.5 were utilized. Following ANC H∞ 28 (EOG-denoised) EEG signals were obtained. For each EEG session data, frequency peaks from (1-15 Hz) of gravity-compensated acceleration signals were identified and removed from the 28 EEG channels via motion artifact removal. Motion artifact denoised EEG data was further cleaned using line noise removal (60 Hz), with Cleanline, Zapline, EEGLAB plug-ins, or Notch filter (60 Hz). In the Artifact Subspace Reconstruction (ASR) algorithm (EEGLAB), the parameter κ = 15 was applied.

### Data Records

All published data are de-identified and subjects gave written consent for their data to be openly shared on a credible public data repository. All data files are available at FigShare (https://figshare.com/projects/The_Slowest_Wave/180343) and have been made available under the terms of Attribution 4.0 International Creative Commons License. The data are archived in a single file set and organized with the following naming convention:

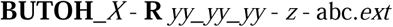

Where *X* is the subject letter (A, B, C, D, and E), *yy_yy_yy* is the date of recording, *z* is the event type (control, rehearsal, calibration), *abc* is the data type (eeg, chanlocs, impedance, opal), .*ext* is the file format (.mat, .set, .h5), and the bold text is fixed for all files.

chanlocs_BoD_mod_32.mat

The chanlocs_BoD_mod_32.mat is a .mat file that contains the EEG electrode locations in 3D space according to the modified 10-20 channel location standard. The file includes the electrode channel labels, channel reference, Cartesian/polar coordinates for each electrode (x, y, z, θ, Φ, r, spherical θ, spherical Φ, and spherical r), channel type and channel number.

-impedance.mat

The -impedance.mat file contains a structure with two fields (variable name: impedance) Impedance.channel: Channel names, including 28 channel EEG, 4-channel EOG, ground and reference

Impedance.value: Impedance values in kΩ

-eeg.set/eeg.mat

The EEG data stored in eeg.set are raw and unfiltered with the four EOG channels.

**Figures**

## Declarations

### Ethics approval and consent to participate

We have complied with all relevant ethical regulations. The study was approved by the institutional review board at the University of Houston, Texas, USA. Informed consent was obtained from all participants.

### Consent for publication

All participants have consented on publishing this manuscript, including on publishing identifying images and videos or other details that compromise anonymity.

### Availability of data and materials

The datasets generated during the current study are available in the Figshare repository (under the terms of Attribution 4.0 International Creative Commons License), the music in Soundcloud, and the dance videos in Vimeo (video links: 1, 2, 3). The underlying code for this study is available in the Figshare repository.

### Competing interests

The authors declare that they have no competing interests.

## Funding

CT wishes to thank The Rockefeller University, Hunter College (City University of New York), the Center for the Ballet and the Arts (New York University), and Gibney Dance.

VG wishes to thank the New York Butoh Institute, the Mellon Foundation, Triskelion Arts, Gibney Dance, Hector Perez and the House of Hallucination, The New York Department of Cultural Affairs, New York Council on the Arts, and the National Endowment for the Arts.

SP wishes to thank The Rockefeller University, the Center for the Ballet and the Arts (New York University), and Gibney Dance.

JCV wishes to thank the NSF IUCRC BRAIN University of Houston Site Award # 2137255, the BRAIN Student Group, and the Noninvasive Brain-Machine Interface Systems Laboratory.

MARM wishes to thank the Associate Dean of Research Office, and the Mechatronics Department at the School of Engineering and Sciences, Tecnológico de Monterrey, and the Noninvasive Brain-Machine Interface Systems Laboratory.

## Authors’ contributions

CT, SP, JCV and VG conceived the study. CT, SP, and JCV designed and co-supervised the study. DH, ET, MRM, and CT ran the mobile EEG recording under supervision of JCV. BK ran the artistic BCI. CT and LB generated Figures 1c and 4b. AMS and CT created Figures 1b, 2 and 6. DH and CT generated Figures 1a, 3 and 5b. VG was the artistic director, choreographer, and dancer of the butoh choreography. CT led the writing of the manuscript, with contributions by SP, JCV, MRM, DH, and BK, while all authors read and gave feedback to the final version of the manuscript.

## Acknowledgments

We would like to acknowledge Michelle Patrick-Krueger, Program Manager of the IUCRC BRAIN at the time of the event for her logistical support.

## Notes

### Competing Interest Statement

The authors have declared no competing interest.

### Summary of Updates

This version of the manuscript has been revised to update in detail how we technically validated all our EEG procedures (e.g., via impedance value monitoring) and how we minimized potential artifacts in our recordings (e.g., via electrooculography and inertial measurement units). We also describe the engineering details and hardware that enabled us to achieve synchronization between signals recorded in different sampling frequencies, and a signal preprocessing and denoising pipeline that we have used to re-sample our data and remove power line noise. As our experiment culminated in a live performance, where we generated a real-time visualization of the dancers' interbrain synchrony on a screen via an artistic brain-computer interface, we outline all the methodology (e.g., filtering, time-windows, equation) we used for online bispectrum estimations.

